# Differential inhibition of GABA release from mouse hippocampal interneuron subtypes by the volatile anesthetic isoflurane

**DOI:** 10.1101/2020.10.01.308460

**Authors:** Iris A. Speigel, Hugh C. Hemmings

## Abstract

General anesthesia is critical to modern medicine and animal research, but the cellular and molecular actions of general anesthetics on the central nervous system remain poorly understood. Volatile anesthetics such as isoflurane disrupt synaptic transmission and inhibit synaptic vesicle release in a neurotransmitter-selective manner. For example, GABA release from interneurons is less sensitive to isoflurane inhibition than are glutamate or dopamine release. Hippocampal and cortical interneuron subpopulations have diverse neurophysiological and synaptic properties, and their individual subtype-specific responses to isoflurane are unknown. We used live-cell optical imaging of exocytosis using fluorescent biosensors expressed in transgenic mouse hippocampal neuron cultures to delineate interneuron subtype-specific effects of isoflurane on synaptic vesicle exocytosis. We found that a clinically relevant concentration of isoflurane (0.5 mM) differentially modulated action potential-mediated exocytosis from GABAergic interneurons: parvalbumin-expressing interneurons were inhibited to 83.1±11.7% of control, whereas somatostatin-expressing and interneurons glutamatergic neurons were inhibited to 58.6±13.3% and 64.5±8.5% of control, respectively. The role of presynaptic voltage-gated sodium channel (Na_v_) subtype expression in determining isoflurane sensitivity was probed by overexpression or knockdown of specific Na_v_ subtypes, which have distinct sensitivities to isoflurane and are differentially expressed between glutamatergic and GABAergic neurons. We found that the sensitivity of exocytosis to isoflurane was determined by the relative expression of Na_v_1.1 (associated with lower sensitivity) and Na_v_1.6 (associated with higher sensitivity). Thus the selective effects of isoflurane on synaptic vesicle exocytosis from hippocampal interneuron subtypes is determined by synaptic diversity in the relative expression of Na_v_1.1 and Na_v_1.6.

**Significance statement:** The volatile anesthetic isoflurane inhibits hippocampal GABAergic interneuron synaptic vesicle exocytosis with differences in potency between interneuron subtypes. This neuron subtype-specific pharmacology derives in part from synaptic diversity in the expression of presynaptic voltage-gated sodium channels that have different sensitivities to anesthetic modulation of channel function. GABAergic interneurons are generally more resistant to the presynaptic effects of isoflurane owing to predominant Na_v_1.1 expression, whereas glutamatergic neurons are more sensitive owing to predominant Na_v_1.6 expression, which supports heterogenous pharmacologic effects on specific neural circuits.

## Introduction

General anesthetics produce both therapeutic and deleterious neurological effects, including postoperative neurocognitive dysfunction, delirium, and developmental neurotoxicity. Despite routine clinical use for over 170 years, as well as important roles in experimental neuroscience, the mechanisms by which these drugs disrupt neural function to cause loss of consciousness, amnesia, and immobility are not fully understand (1).

General anesthetics have prominent actions on synaptic transmission through multiple presynaptic and postsynaptic sites of action. Progress in studying molecular anesthetic mechanisms has focused largely on postsynaptic modulation of neurotransmitter receptors, including potentiation of inhibitory GABA_A_ receptors and inhibition of excitatory NMDA receptors. Far less is known about their presynaptic actions due in part to the technical challenges in distinguishing presynaptic and postsynaptic drug effects in intact synapses. Volatile general anesthetics like isoflurane inhibit synaptic vesicle (SV) exocytosis and neurotransmitter release from isolated nerve terminals and from primary neurons (2, 3). Glutamatergic synaptic transmission is inhibited with greater potency than GABAergic transmission, generating net depression of central nervous system function. The molecular basis for this synapse-specific anesthetic pharmacology is unknown. Identifying the presynaptic targets that underlie transmitter-specific differences in anesthetic potency is critical for mechanism-driven optimization of anesthetic agent pharmacology.

Exocytosis-mediated neurotransmitter release is determined primarily by the amount of Ca^2+^ entering the synaptic terminal (4). Isoflurane decreases presynaptic Ca^2+^ entry with no apparent effect on neuronal Ca^2+^/exocytosis coupling, implicating targets that control membrane depolarization and Ca^2+^ entry. Changes in presynaptic Ca^2+^ depend on depolarization by the invading action potential, which is shaped by presynaptic receptors and Na^+^, Ca^2+^ and K^+^ ion channels, many of which are affected by anesthetics.

Excitatory and inhibitory neurons express different complements of presynaptic ion channels and receptors that regulate synaptic function. Hippocampal and cortical interneurons are comprised of functionally diverse subtypes with characteristic neurophysiological properties and differential voltage-gated ion channel expression (5). Hippocampal GABAergic interneurons expressing parvalbumin (PV^+^) are distinguished by their high axonal Na_v_1.1 expression compared to glutamatergic neurons, which enables a fast-spiking phenotype (6). Interneuron subtypes also differ in their postsynaptic targets and neural circuit functions; PV^+^ and somatostatin (SST^+^)- expressing interneurons target pyramidal cells and suppress glutamate release, whereas vasoactive intestinal peptide (VIP^+^)-expressing interneurons target other interneurons and contribute to circuit disinhibition (5).

While the molecular machinery mediating synaptic vesicle exocytosis is remarkably conserved between synapses, volatile anesthetics like isoflurane differentially depress neurotransmitter release, with GABA release markedly less inhibited than glutamate release (7). The molecular bases of this transmitter-specific selectivity are critical to understanding the effects of anesthetics on neuronal function. Given the critical roles of various hippocampal interneuron subtypes in learning and memory (5), we addressed the relative anesthetic sensitivities of specific interneuron subtypes, which is critical to our conceptual understanding of anesthetic mechanisms at both the cellular and network levels. We hypothesized that the volatile anesthetic isoflurane has selective pharmacological actions of GABAergic hippocampal interneuron subtypes based on their presynaptic molecular diversity. We investigated interneuron subtype-specific anesthetic actions using cultured hippocampal neurons derived from transgenic mice in which GABAergic interneuron subtypes are identified by expression of genetically encoded fluorescent markers.

## Results

### Isoflurane has heterogeneous effects on synaptic vesicle exocytosis from GABAergic interneurons

Anesthetic actions on SV exocytosis from GABAergic neurons was determined from primary hippocampal neuron cultures derived from GAD2-Cre::Ai14-tdTomato transgenic mice, a cross of the GAD2-Cre and the Ai14 Cre-dependent reporter strain. The GAD2 gene encodes the GABA synthesizing enzyme glutamic acid decarboxylase (GAD65), a specific marker for GABAergic neurons (8), such that Cre-dependent expression of the red fluorescent protein tdTomato genetically labels GABAergic (GAD2^+^) axons and terminals (**Figure 1A**)(9). This GAD2Cre line has been validated by GAD65 immunohistochemistry to target cortical and hippocampal GABAergic neurons with nearly complete specificity and efficiency (9-11); immunostain analysis of primary hippocampal neuron cultures from this GAD2-Cre crossed with another reporter line shows that genetically labeled GAD2^+^ neurons have no overlap with CAMKII, a glutamatergic neuron marker (12).

**Figure 1.**
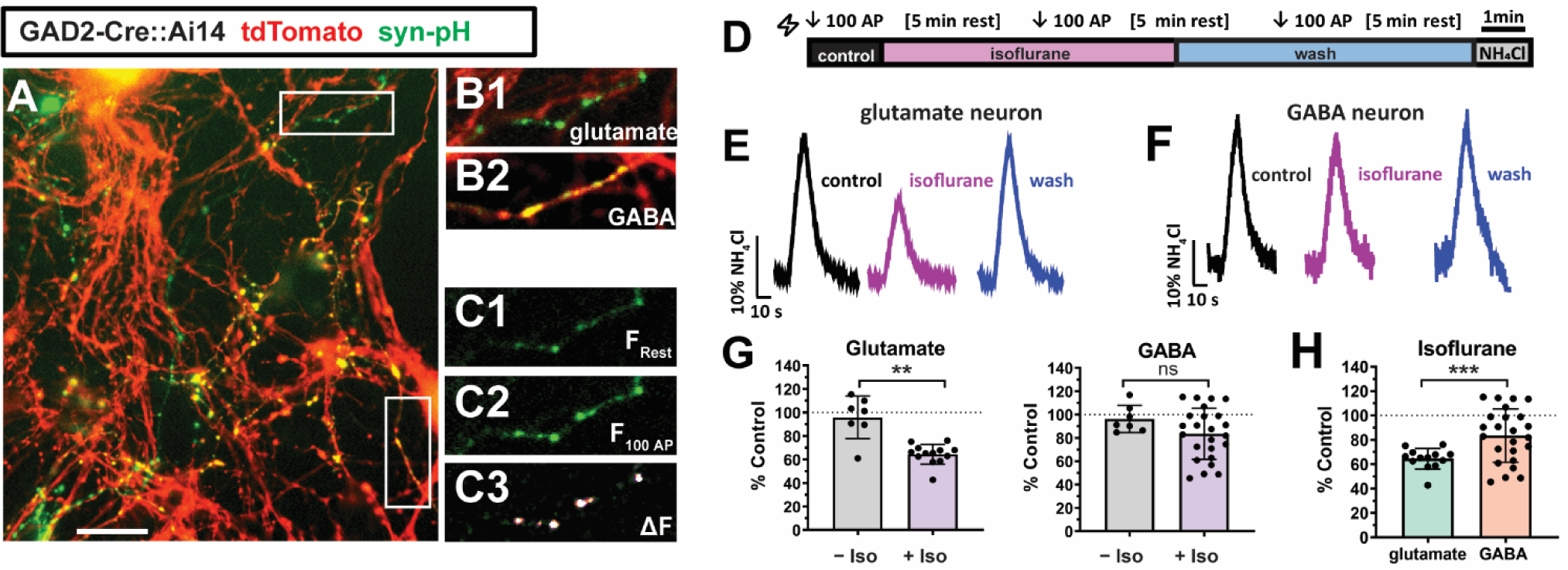
Synaptic vesicle exocytosis from mouse hippocampal interneurons shows reduced inhibition by isoflurane. **A**. GAD2^-^ and GAD2^+^ (putative glutamatergic and GABAergic, respectively) neurons are distinguishable in cultures of GAD2-Cre::Ai14 transgenic mouse hippocampal neurons based on cell-specific td-Tomato expression. Synaptophysion-pHluorin (syn-pH)-transfected glutamatergic (GAD2^-^) and GABAergic (GAD2^+^) neuron processes are either green (syn-pH^+^) or yellow (red[td-Tomato]/green [syn-pH] overlap), respectively. Boutons arising from the same neuron can be identified as a “string of pearls” or isolated lengths of axon with *en passant* boutons identified by syn-pH fluorescence. Scale bar = 10 µm. **B1-2**. Glutamatergic (Glutamate/GAD2^-^) and GABAergic (GABA/GAD2^+^) syn-pH-expressing boutons; figures are from insets of 1A, and the lower inset representing GABAergic boutons is rotated 90 degrees. Length of figure = 15 µm. **C**. Electrical activity-driven changes in syn-pH fluorescence reflect synaptic vesicle (SV) alkalization upon exocytosis and re-acidification following endocytosis. F_Rest_ (**C1**) represents the resting fluorescence prior to 100 action potentials (AP; F100 AP; **C2**), and ΔF (**C3**) represents the difference. **D**. Live-cell imaging protocol to determine isoflurane effects on SV exocytosis. Control, isoflurane (0.5 mM; ∼1.4 MAC) and wash-in control buffer (Tyrode’s solution, pH 7.4, plus 2 mM Ca^2+^, 10 µM CNQX, and 50 µM APV) continuously perfused at ∼1 ml/min. SV exocytosis was elicited by electrical stimulation (100 AP at 10 Hz, 30 mA/ms, 10 V/cm^2^ field), with 5 min rest periods. NH_4_Cl perfusion identifies the total pool of releasable vesicles. **E-F**. Representative traces from glutamatergic or GABAergic neurons showing SV exocytosis evoked by 100 AP for control and isoflurane perfusion. Control stimulations reached 24% and 36% respectively of the total releasable pool of SVs revealed by NH_4_Cl perfusion. Isoflurane inhibition of 100 AP-evoked SV exocytosis was reversible with washout by control buffer. **G**. Mean inhibition of ΔF/F_0_ for glutamatergic and GABAergic neurons by isoflurane (+ Iso) shown as % control, compared to control recordings without isoflurane (− Iso). Isoflurane inhibited SV exocytosis in glutamate neurons compared to control (-Iso; ** p=0.0028). **H)** Comparison of isoflurane inhibition of SV exocytosis in glutamatergic *vs* GABAergic neurons. Isoflurane (0.5 mM) inhibited SV exocytosis more in glutamatergic than in GABAergic hippocampal neurons (glutamate (GAD2^−^): 64±8% of control, n=13; GABA (GAD2^+^): 84±22% control, n=24, *** p=0.0006, unpaired t-test). GABAergic neurons showed variability in sensitivity to isoflurane, with some showing greater SV exocytosis in the presence of isoflurane compared with control.

To measure SV exocytosis, cultures were transfected with the optical biosensor synaptophysin-pHluorin (syn-pH) at low efficiency via calcium phosphate-mediated gene transfer. Syn-pH is a chimera of the SV-associated protein synaptophysin and a pH-sensitive eGFP (pHluorin) that reports exocytosis upon exposure of the acidic SV interior to the extracellular (neutral pH) (13). GABAergic boutons can be readily identified by co-expression of syn-pH and tdTomato since syn-pH expressing terminals are visible due to high basal expression on the plasma membrane (**Figure 1B2**). Transfected boutons negative for tdTomato are identified as GAD2^−^ boutons arising from putative glutamatergic neurons (**Figure 1B1)**. Given the low efficiency of calcium phosphate mediated transfection, syn-pH expression within a single field of view could be attributed to a single interneuron. Preliminary studies to assess transfection efficiency using eGFP showed an average 0.75 interneurons were transfected per coverslip, for a GAD2^+^ transfection efficiency of ∼0.157%.

A brief train of 100 action potentials (AP) was generated by electrical stimulation at 10 Hz to evoke SV exocytosis. Stimulation with 100 APs evoked a rapid rise in syn-pH fluorescence, which was quantified in 2 μm regions of interest (ROI) set to include responsive boutons (**Figure 1C**). The experimental paradigm for recording, stimulation, and solution exchange is illustrated in **Figure 1D**. Exocytosis was measured with perfusion of control buffer before and after perfusion of 0.5 mM isoflurane, which corresponds to a clinically relevant dose of ∼1.4 times the minimum alveolar concentration (MAC) adjusting for species and temperature (14). Isoflurane concentration was verified by gas chromatography of perfusate to be 0.50±0.039 mM. Representative traces from a glutamatergic or GABAergic neuron are shown in **Figure 1E and F**, respectively. In both cell types isoflurane inhibition of SV exocytosis was rapidly reversible upon washout which indicates that the observed inhibition was not due to deteriorating cell conditions.

Isoflurane inhibited SV exocytosis, expressed as percentage of control, in glutamatergic neurons to a greater extent than in GABAergic neurons (GAD2^-^: 64.5±8.5%, n=13; GAD2^+^: 83.7±22.0%, n=24, p=0.0006, Welch’s t-test) (**Figure 1H**). The population averages of GAD2^+^ (GABAergic) and GAD2^−^ (glutamatergic) neurons observed was consistent with previous pHluorin-based live-cell imaging studies and neurochemical assays (3, 15). However, the pool of recorded GAD2^+^ neurons showed variation in the isoflurane effect at the single-cell level. Although most neurons linshowed inhibition by isoflurane, in 20% (5/24) of recorded GAD2^+^ neurons SV exocytosis was potentiated by isoflurane. This cell-type specific difference was not reflected in sham recordings in which isoflurane was omitted in order to test for reproducibility of the 100 AP-driven syn-pH ΔF/F_0_ signal over three serial stimulations in an identical recording paradigm (**Figures 1E-G** and **Supplemental Figure 1**). **Supplementary Figure 2** shows a histogram of GAD2^+^ SV exocytosis modulation by isoflurane. We hypothesize that this variance reflects functional diversity of hippocampal interneurons with commensurate pharmacological diversity in sensitivity to the volatile anesthetic isoflurane.

**Figure 2.**
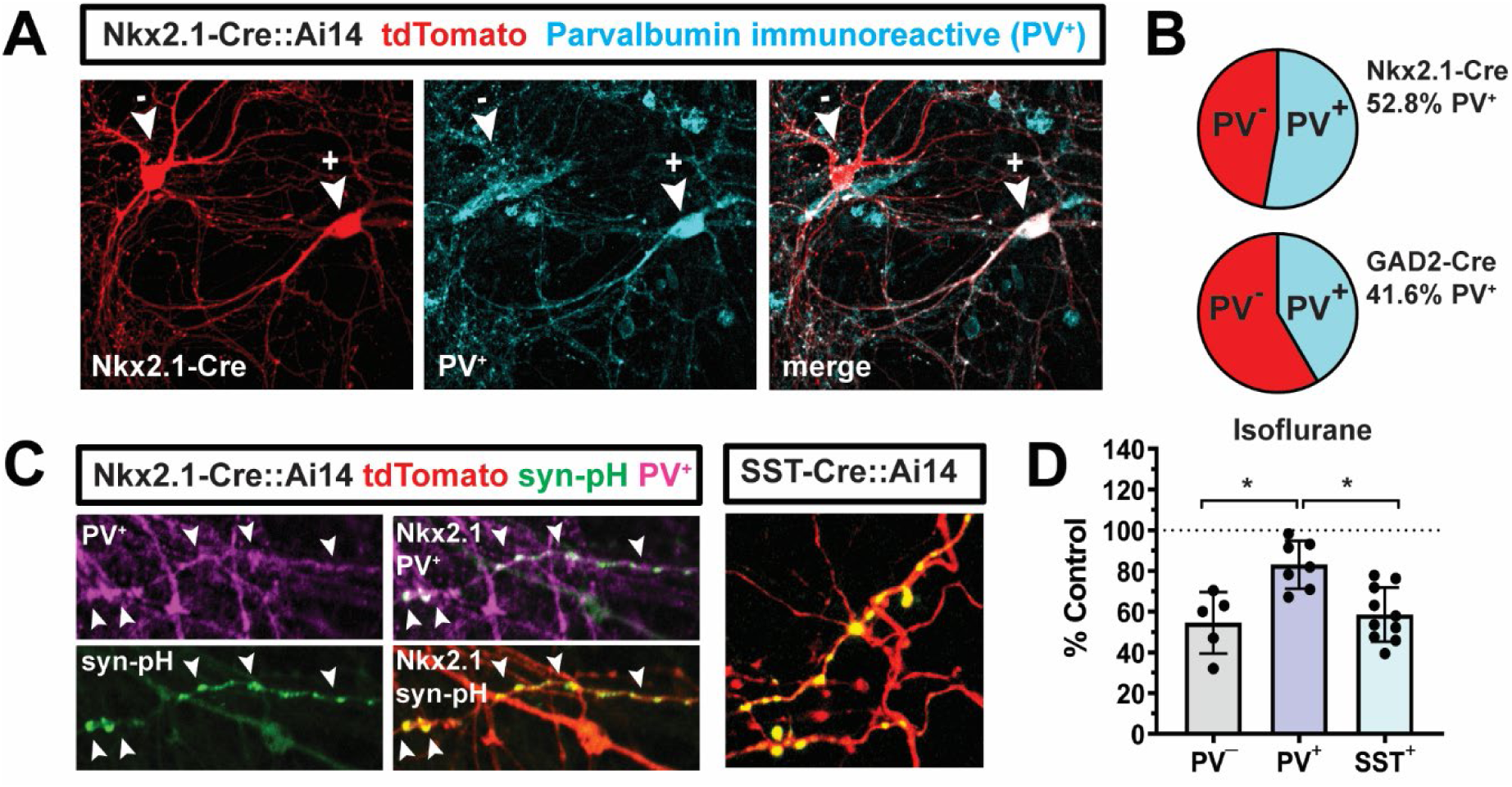
Isoflurane differentially inhibits SV exocytosis from parvalbumin- and somatostatin-expressing interneurons. **A**. Hippocampal interneurons from Nkx2.1Cre::Ai14 transgenic mice include tdTomato-expressing PV immunoreactive (PV^+^; + arrowhead) and non-immunoreactive interneurons (− arrowhead). Scale bar = 10 µm. **B**. PV^+^ neurons represent 53% of Nkx2.1^+^ neurons (n=57), and 41% of GAD2^+^ neurons (n=549) in Nkx2.1-Cre::Ai14 and GAD2-Cre::Ai14 transgenic cultures, respectively. **C**. Left panel, Nkx2.1^+^ boutons expressing transfected syn-pH can be identified and evaluated as PV^+^ or PV^−^ by retrospective immunocytochemistry. Arrowheads indicate boutons triple-positive for PV, Nkx2.1/tdTomato, and syn-pH. Figure length = 24 µm. Right panel from SST::Ai14 culture shows SST^+^ axon and boutons (red) transfected with syn-pH (yellow). Figure length = 41 µm. **D**. Isoflurane inhibited SV exocytosis from Nkx2.1^+^/PV^−^ neurons (left) and Nkx2.1^+^/PV^+^ neurons (middle) and SST^+^ neurons (right). There was greater inhibition of PV^−^ and SST^+^ neurons compared to PV^+^ neurons (PV^−^ = 55±15% of control, n=5; PV^+^ = 83±12% of control, n=7; SST^+^ = 59±13% control, n=10) (PV^−^ *vs*. PV^+^, ** p=0.0044; PV^+^ *vs*. SST^+^, ** p=0.0037; all ANOVA with Tukey’s *post hoc* test).

### Differential effects of isoflurane on hippocampal Interneuron subtypes

We examined GABAergic interneuron subtype-specific anesthetic actions on SV exocytosis using cultures from transgenic mice with genetically labelled interneuron subpopulations. We focused on parvalbumin- and somatostatin-expressing interneurons (PV^+^ and SST^+^) due to their relative abundance and importance in cortical and hippocampal circuit function (5).

Immunocytochemical analysis of GAD2-Cre::Ai14 cultures at DIV 18, the average age used for imaging experiments, revealed that 41.6% of GAD2^+^ neurons expressed parvalbumin (PV) (**Figure 2B**). To target PV^+^ interneurons for SV exocytosis measurements, we used Nkx2.1-Cre::Ai14 hippocampal cultures in combination with retrospective immunostaining for PV. The Nkx2.1 progenitor label was chosen as a marker for PV^+^ neurons because PV expression has late developmental onset, beginning during the second postnatal week and stabilizing after the third postnatal week (16). PV-Cre::Ai14 and PV-tdTomato mice did not show neuronal reporter protein expression <DIV21, requiring an earlier genetic label for SV exocytosis experiments. Nkx2.1-Cre targets the neural progenitors that give rise to PV^+^ and SST^+^ neurons, enabling PV^+^ cells to be labeled prior to active transcription of PV^+^ (17). At DIV 18, Nkx2.1-Cre::Ai14 cultures showed that 52.8% of Nkx2.1^+^ cells were PV-immunoreactive (**Figure 2A-B**).

Cultures from NKX2.1-Cre::Ai14 mouse hippocampus were transfected with syn-pH for measurement of SV exocytosis, and fixed for *post hoc* PV immunofluorescence (**Figure 2C, left**). Isoflurane inhibited SV exocytosis in PV^+^ (PV-immunoreactive) neurons to 83.1±11.7% of control, whereas PV^−^ (PV-negative) neurons were inhibited to 54.6±15.1% of control (p=0.0044, ANOVA with Tukey’s *post hoc*) (**Figure 2D**). Because Nkx2.1^+^ neurons include the mutually exclusive PV and SST interneuron subpopulations, we hypothesized that the PV^-^ group included SST^+^ interneurons and/or PV neurons that were weakly immunoreactive due to delayed PV protein expression.

For verification of the distinct sensitivity of SST^+^ interneurons to isoflurane inhibition of SV exocytosis, we measured SV exocytosis in SST-Cre::Ai14 cultures. SST is expressed prenatally (18), and SST^+^ neurons genetically labeled with tdTomato were identified *in vitro* from hippocampal neuron cultures (**Figure 2C, right**). Isoflurane inhibited SV exocytosis in SST^+^ neurons to 58.6±13.3% of control, significantly greater than the effect on PV^+^ neurons (p=0.0037, ANOVA with Tukey’s *post hoc*) (**Figure 2D, right**). Isoflurane inhibited SV exocytosis from SST^+^ neurons to a similar extent as glutamatergic (GAD2^−^) neurons, which was significantly greater than for the GABAergic (GAD2^+^) population average (SST^+^ *vs* GAD2^−^, p=0.2078; SST^+^ *vs* GAD2^+^, p=0.0040; both t-test).

### Overexpression of Na_v_1.1 and Na_v_1.6 alters sensitivity of SV exocytosis to isoflurane

Voltage-gated Na^+^ channels (Na_v_) control presynaptic excitability, voltage-gated Ca^2+^ channel (Ca_v_) activation, Ca^2+^ influx, and Ca^2+^-coupled neurotransmitter release (4, 19). Neuronal Na_v_ subtypes are differentially inhibited by volatile anesthetics in a channel subtype-selective manner (Na_v_1.1 < Na_v_1.2 ≅ Na_v_1.6), such that differential expression between interneuron subtypes could lead to differences in isoflurane effects on SV exocytosis (20). Na_v_1.1 is enriched in interneurons and is the predominant Na_v_ subtype expressed in GABAergic boutons, whereas Na_v_1.6 is the dominant subtype expressed in glutamatergic pyramidal neurons (21). Such synaptic heterogeneity manifest as differences in Na_v_1.1 expression and subcellular compartmentalization between interneuron subtypes could explain differential pharmacological effects between interneurons (22).

We hypothesized that relatively greater expression of Na_v_1.1 in GABAergic boutons leads to reduced sensitivity to isoflurane, whereas relatively greater expression Na_v_1.6 in glutamatergic boutons leads to increased sensitivity to isoflurane. We tested this hypothesis by overexpression of Na_v_1.1 and Na_v_1.6 with syn-pH based imaging of SV exocytosis. Anesthetic effects determined by specific Na_v_ subtypes and confirmation that overexpressed channels functionally integrated into the presynaptic machinery and contributed to neurotransmitter release was facilitated by employing Na_v_ mutants rendered resistant to tetrodotoxin (TTX-R) by point mutations in each Na_v_ subtype: Na_v_1.1_R_ [Na_v_1.1 F383S] and Na_v_1.6_R_ [Na_v_1.6 Y371S] (23, 24). In the presence of 300 nM TTX, SV exocytosis in cultured hippocampal neurons without co-transfection with TTX-R Na_v_s was completely abolished. Transfection of either Na_v_1.1_R_ or Na_v_1.6_R_ rescued action potential-evoked SV exocytosis in the presence of TTX (**Figure 3A-B**). Immunocytochemical labelling of Na_v_1.1 and Na_v_1.6 shows increased expression following transfection of Na_v_1.1_R_ or Na_v_1.1_R_, confirming that the mutant channels can be synthesized and trafficked to axons (**Supplemental Figure 4**)

**Figure 3:**
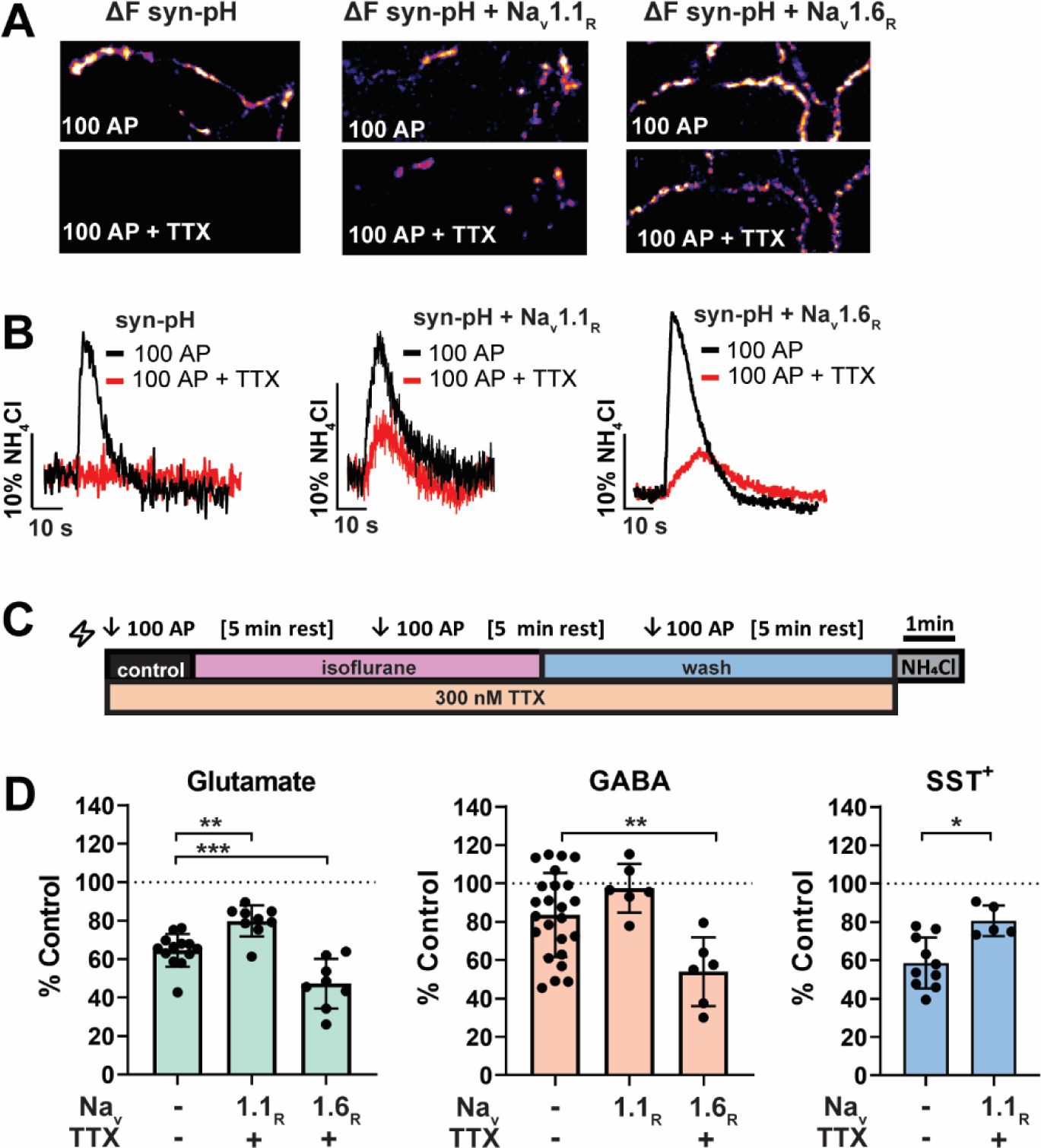
Expression of tetrodotoxin-resistant Na_v_1.1_R_ or Na_v_1.6_R_ identifies selective roles in isoflurane inhibition of SV exocytosis. **A**. Co-transfection of syn-pH with the tetrodotoxin resistant Na_v_ mutants hNa_v_1.1_R_ or mNa_v_1.6_R_ rescues SV exocytosis from inhibition by tetrodotoxin (TTX, 300 nM), which completely inhibited exocytosis mediated by endogenous channels. Recordings of Na_v_1.1_R_ or Na_v_1.6_R_ mediated exocytosis in the presence of TTX were performed using Tyrode’s solution adjusted to 4 mM Ca^2+^, 0 mM Mg^2+^. Figure lengths = 24 µm. **B**. Representative traces showing SV exocytosis in the presence of TTX (red), which abolished SV exocytosis in neurons expressing only syn-pH (left), but not in neurons co-transfected with Na_v_1.1_R_ or Na_v_1.6_R_ (middle, right). **C**. Live-cell imaging protocol to isolate Na_v_1.1_R_-mediated or Na_v_1.6_R_-mediated SV exocytosis and measure selective isoflurane effects. Control, isoflurane (0.5 mM) and wash solutions are prepared in Tyrode’s buffer plus 4 mM Ca^2+^, 300 nM TTX, 10 µM CNQX, and 50 µM APV. **D**. Isoflurane effect on SV exocytosis displayed as percent of control, taken from neurons transfected with Na_v_1.1_R_ or Na_v_1.6_R_. Degree of isoflurane inhibition is significantly different compared to neurons transfected only with syn-pH (isoflurane effects measured in 2 mM Ca^2+^ Tyrode’s solution without TTX). **Left, Glutamatergic neurons (GAD2**^−^): endogenous Na_v_ *vs*. Na_v_1.1_R_, ** p=0.0023, endogenous Na_v_ *vs*. Na_v_1.6_R_ *** p=0.0009, ANOVA with Dunnett’s *post hoc* test. endogenous Na_v_, n=13, Na_v_1.1_R_, n=7, Na_v_1.6_R_, n=8. **Middle, GABAergic neurons (GAD2**^**+**^): endogenous Na_v_ *vs*. Na_v_1.1_R_, p=0.273, endogenous Na_v_ *vs*. Na_v_1.6_R_, p=0.0068, ANOVA with Dunnett’s *post hoc* test. Endogenous Na_v_, n=24, Na_v_1.1_R_, p=6, Na_v_1.6_R_, n=6. **Right, somatostatin neurons (SST**^**+**^**/GAD2**^**+**^): endogenous Na_v_ *vs*. Na_v_1.1_R_, p=0.011, t-test. Endogenous Na_v_, n=10; Na_v_1.1_R_, n=5.

**Figure 4:**
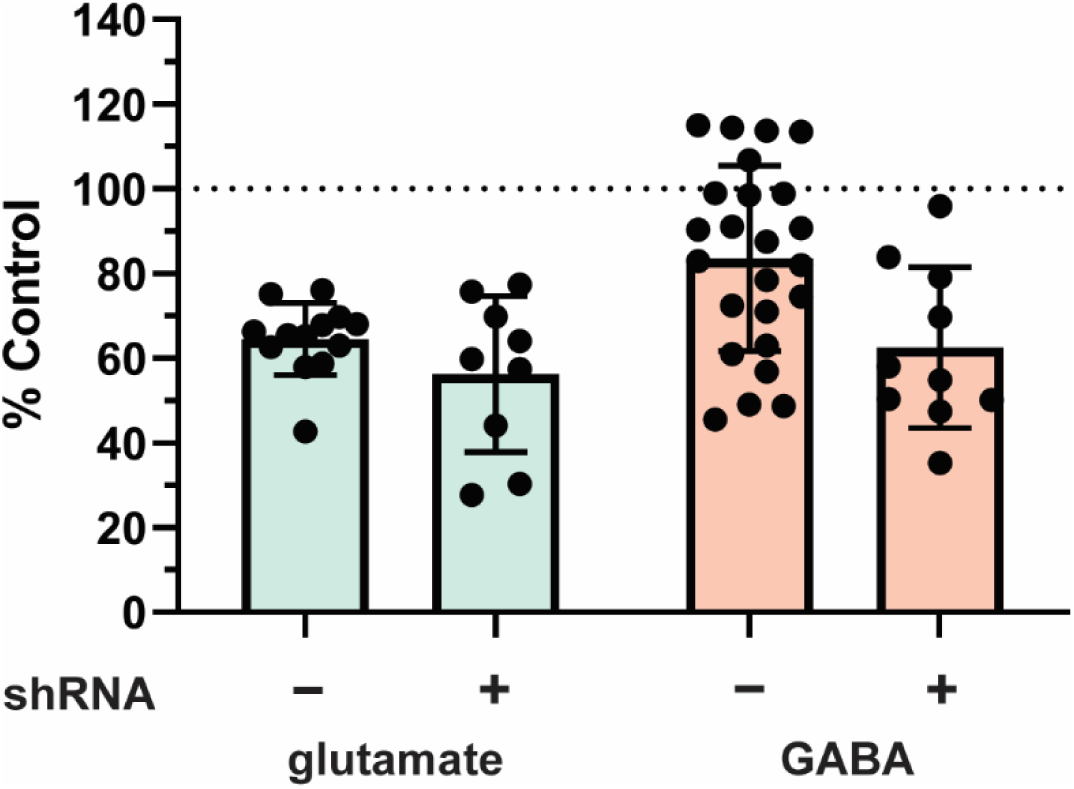
Knockdown of Na_v_1.1 increases inhibition of SV exocytosis by isoflurane in GAD2^+^ interneurons. Isoflurane inhibition of SV exocytosis in GABAergic/GAD2^**+**^ neurons co-transfected with syn-pH plus Na_v_1.1 shRNA compared to GAD2^**+**^ neurons transfected only with syn-pH (+ shRNA = 62%±19 *vs*. – shRNA = 84±22%, p=0.013, t-test n=10). Isoflurane inhibition in glutamatergic/GAD^−^ neurons co-transfected with Na_v_1.1 shRNA was not significantly different from neurons without (+ shRNA = 56%±18 *vs*. – shRNA = 64±8, p=0.17, t-test, n=9).

We analyzed the effects of isoflurane on Na_v_ subtype-specific exocytosis by co-transfection of syn-pH with Na_v_1.1_R_ or Na_v_1.6_R_ with measurement of SV exocytosis in the presence of 300 nM TTX to block native Na_v_ (**Figure 3C**). Inhibition by isoflurane was quantified as a percent of control in each group compared to isoflurane inhibition previously measured for native Na_v_ in neurons in the absence of TTX (**Figure 3D, left and middle**). In glutamatergic neurons from GAD2::Ai14 cultures, isoflurane showed diminished inhibition in Nav1.1_R_-expressing neurons compared to neurons lacking mutant Na_v_, and increased inhibition in Na_v_1.6_R_-expressing neurons (GAD2^−^ no mutant Na_v_: 64.5±8.5%; Na_v_1.1_R_: 79.8±8.1%; Na_v_1.6_R_: 47.1±12.9%; no mutant Na_v_ *vs*. Na_v_1.1_R_ p=0.0023, no mutant Na_v_ *vs*. Na_v_1.6_R_ p=0.0009, ANOVA with Dunnett’s *post hoc* test). In GAD2^+^ neurons with Na_v_1.1_R_ expression, isoflurane inhibition was not different, whereas Na_v_1.6 _R_ expressing neurons showed greater inhibition by isoflurane (GAD2^+^ no mutant Na_v_: 83.5±21.9%, Na_v_1.1_R_: 98.38±14.0%, Na_v_1.6_R_ 53.9±17.9%, no mutant Na_v_ *vs*. Na_v_1.1_R_ p=0.273, NT *vs*. Na_v_1.6_R_ p=0.0068, ANOVA with Dunnett’s *post hoc* test).

We hypothesized that SST^+^ interneurons are more sensitive to isoflurane inhibition than PV^+^ interneurons due to lower expression of axonal Na_v_1.1 (22, 25). Using Na_v_1.1_R_ expression in SST-Cre::Ai14 cultures, SST^+^ neurons expressing Na_v_1.1_R_ showed less inhibition by isoflurane compared with those not expressing Na_v_1.1_R_ (SST^+^: no mutant Na_v_ 58.6±13.3%; Na_v_1.1_R_ 78.0%±6.642, p=0.018, t-test) (**Figure 3D, right**).

### Knockdown of Na_v_1.1 in GABAergic neurons increases anesthetic sensitivity of exocytosis

Further evidence for the role of Na_v_1.1 in GABAergic interneuron subtype-specific anesthetic actions was obtained using short hairpin RNA (shRNA) to knockdown endogenous Na_v_1.1 in cultured hippocampal neurons. Following identification and validation of an effective Na_v_1.1 shRNA construct (**Supplementary Figure 5**), we co-transfected GAD2-Cre::Ai14 transgenic hippocampal neuron cultures with Na_v_1.1 shRNA and syn-pH. In comparing isoflurane inhibition of SV exocytosis in control *vs*. shRNA-treated Na_v_1.1 neurons (**Figure 4**), shRNA-mediated knockdown of Na_v_1.1 increased inhibition by isoflurane of SV exocytosis compared to untreated GABAergic/GAD2^+^ neurons, but had no effect on glutamatergic/GAD2^−^ neurons (GAD2^+^: + shRNA 62.5%±19.0 *vs*. – shRNA 83.7±22.0%, p=0.013, t-test; glutamatergic/GAD2^−^ neurons: + shRNA 56.3%±18.4 *vs* – shRNA 64.5±8.5, p=0.17, unpaired t-test). This neuron subtype-specific effect is consistent with greater endogenous Na_v_1.1 expression in GABAergic interneurons (49).

**Figure 5:**
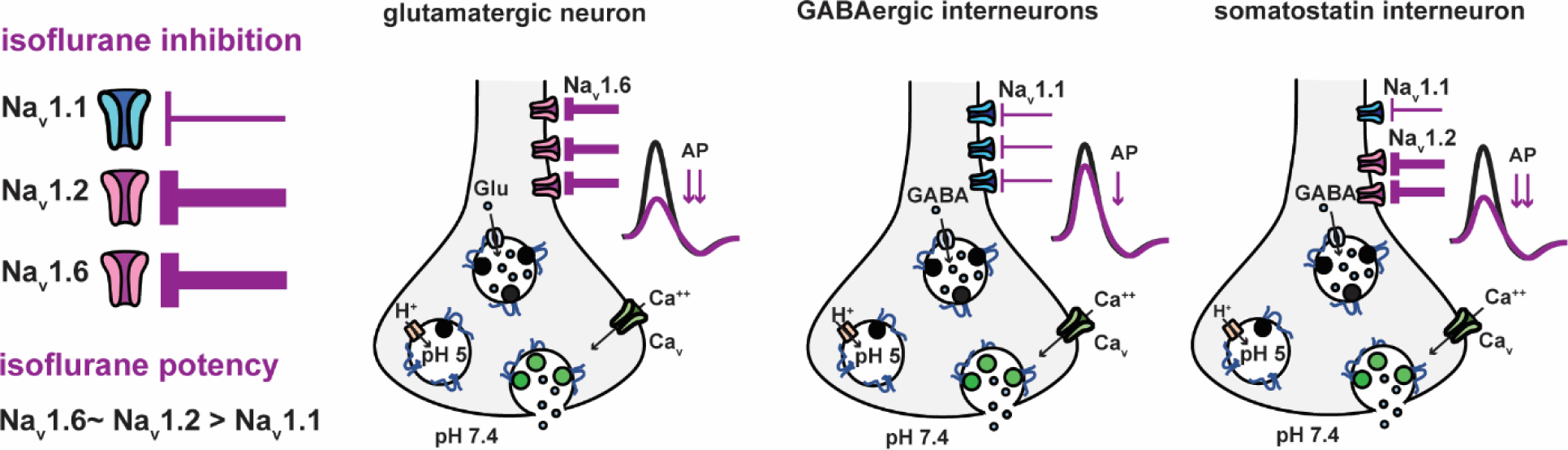
Summary of presynaptic mechanisms contributing to differential inhibition of SV exocytosis by isoflurane. The three major CNS Na_v_ subtypes have distinct sensitivities to isoflurane modulation of channel function, with cell-specific differences in expression influencing cellular responses to isoflurane. **A**. Na_v_1.1 (blue) is relatively resistant to inhibition compared to Na_v_1.2 and Na_v_1.6 (pink) at normal resting membrane potential. **B**. Hippocampal glutamatergic neurons are sensitive to isoflurane depression of the presynaptic action potential due to predominant Na_v_1.6 expression, resulting in reduced terminal depolarization, presynaptic Ca^2+^ entry, and SV exocytosis/transmitter release. GABAergic interneurons are less sensitive to isoflurane due to relatively greater Na_v_1.1 expression. Because PV^+^ interneurons are a major representative GABAergic/GAD2^+^ cell type, isoflurane inhibition of SV exocytosis in GAD2^+^ cells is skewed to reflect the reduced sensitivity of PV^+^ neurons to isoflurane. Somatostatin (SST^+^) interneurons express relatively less Na_v_1.1 and more Na_v_1.2, which has a similar sensitivity to isoflurane as Na_v_1.6, resulting in isoflurane sensitivity comparable to glutamatergic neurons.

## Discussion

We posited that general anesthetics have differential neural circuit effects based on interneuron subtype-specific anesthetic pharmacology as a result of synaptic diversity in the differential expression of synaptic signaling molecules.

### Diverse anesthetic actions on GABAergic neurons

Using GAD2-driven expression of fluorescent markers to identify hippocampal interneurons, we found that specific interneuron subtypes exhibited differences in isoflurane sensitivity. This was not apparent in previous studies of GABAergic boutons identified by a retrospective labelled vGAT antibody uptake method (2) due to the low-throughput nature of this technique. Use of GAD2 transgenic cultures allowed targeted analysis of GAD2^+^ interneurons. Our analysis of SV exocytosis in hippocampal interneurons revealed a surprising range of anesthetic effects: some interneurons had a similar sensitivity as glutamatergic neurons, whereas others were potentiated by isoflurane with increased SV exocytosis. We hypothesized that the effects of isoflurane on SV exocytosis differ depending on the cellular phenotype and synaptic diversity in the hippocampus, as has been observed for other neuronal subtypes in other brain regions (26-28). The potentiation of GABAergic SV exocytosis is consistent with previous neurochemical and electrophysiological findings of anesthetic-induced GABA release (29, 30).

We focused our analysis of anesthetic actions on PV^+^ and SST^+^ interneurons, the most numerous and well-characterized cortical and hippocampal interneuron subpopulations. Although both arise from the Nkx2.1-expressing medial ganglionic eminence embryonic developmental cohort, PV^+^ and SST^+^ interneurons exert distinct forms of GABAergic inhibition. PV^+^ interneurons preferentially synapse on the perisomatic region of pyramidal cells and have fast-spiking bursts that produce robust large-scale inhibition of neural activity (31), while SST^+^ interneurons preferentially synapse on distal dendrites to inhibit specific synaptic inputs locally (5).

Our live-cell imaging analysis of SV exocytosis from identified PV^+^ and SST^+^ interneurons revealed their distinct sensitivities to isoflurane. PV^+^ neurons were inhibited to 78% from control, comparable to the GAD2^+^ population average of 83%. Despite many similarities, SST^+^ interneurons showed greater inhibition by isoflurane (inhibited to 58.6% of control), comparable to glutamatergic neurons (inhibited to 64.5% of control). Immunochemical analysis of our cultures revealed that about half of GAD2^+^ cells were PV^+^, which is consistent with other reports both *in vitro* and *in vivo (16)*. Previous determinations of the potency of isoflurane inhibition of GABA release determined using biochemical and imaging methods represent population averages that are driven lower compared to glutamate release by the PV^+^ interneuron response.

Notably, isoflurane increased SV exocytosis in ∼20% of GAD2^+^ interneurons, consistent with an increase in transmitter release. Because this response was never observed in PV^+^ and SST^+^ interneuron recordings, or in any recordings from NKX2.1^+^ neurons, we hypothesize that potentiation occurred in NKX2.1^-^ interneurons, an embryonic cohort that includes cholecystokinin (CCK)-, vasoactive intestinal peptide (VIP)-, and (CR)-expressing interneurons. VIP^+^ and CR^+^ neurons preferentially target other interneurons for circuit disinhibition (5). Isoflurane potentiation of GABA release is consistent with electrophysiological recordings of CA1 interneurons in which isoflurane increased spontaneous inhibitory postsynaptic current (IPSC) frequency, suggesting potentiated GABAergic input onto these interneurons (32). A potential consequence of this would be anesthetic-induced circuit disinhibition, which might contribute to paradoxical excitation and burst suppression, two unexplained phenomena characterized by spontaneous neural hyperactivity during light or deep anesthesia, respectively (33). Neuronal potentiation by isoflurane is best characterized in the hypothalamus, where circuits common to sleep and anesthesia appear to be targets for hypnotic drug actions. Ventrolateral preoptic non-noradrenergic neurons are depolarized by isoflurane as a result of diminished postsynaptic K^+^ conductance (34). Activity of a mixed GABAergic/glutamatergic cluster of neuroendocrine cells within and near the supraoptic nucleus increases during anesthesia (35). These nuclei are most likely related to anesthetic-induced loss of consciousness, while potentiation of circuits in the hippocampus would be more relevant to amnesia.

### Specific Na_v_ isoform expression determines interneuron subtype-specific pharmacology

Synaptic vesicle exocytosis is supralinearly dependent on presynaptic Ca^2+^ influx through Ca_v_ activated by nerve terminal depolarization. Many of the presynaptic ion channels that contribute to the AP waveform (Na^+^, Ca^2+^, and K^+^ channels) are sensitive to anesthetics such that their pharmacological modification can have significant consequences for transmitter release (36). Multiple Ca_v_ subtypes are inhibited by isoflurane (37), and show differential expression in hippocampal interneurons with P/Q- and N-type channels being the predominant presynaptic Ca_v_ subtypes (38). However since isoflurane inhibits P/Q- and N-type channel-coupled SV exocytosis with comparable potencies (39) we decided to focus on Na_v_, putative presynaptic targets upstream of Ca_v_.

Neuronal Na_v_s contribute to presynaptic depolarization by generating the depolarizing/rising phase of the action potential. Na_v_1.1 is significantly less sensitive to isoflurane than Na_v_1.2 and 1.6 (20); at a physiological resting membrane potential of -70 mV, Na_v_1.1 peak current is resistant to 1.6 MAC isoflurane whereas Na_v_1.2 and Na_v_1.6 are significantly inhibited. Because Na_v_1.1 is enriched in GABAergic interneurons while Na_v_1.6 is more abundant in glutamatergic neurons, we hypothesized that Na_v_1.1 expression in GABAergic boutons determines their reduced sensitivity to isoflurane inhibition of SV exocytosis. Na_v_1.1 overexpression diminished the isoflurane sensitivity of SV exocytosis in glutamatergic but not in GAD2^+^ GABAergic neurons. This is consistent with greater Na_v_1.1 abundance in GAD2^+^ neurons, and the high Na_v_1.1 expression in PV^+^ interneurons, a large subpopulation. Conversely, both Na_v_1.1 knockdown and Na_v_1.6 overexpression enhanced isoflurane sensitivity in GAD2^+^ neurons. Together these results show that the relative expression of Na_v_1.1 *vs* Na_v_1.6 contributes to presynaptic sensitivity to isoflurane, and that altering their relative expression can modulate interneuron sensitivity to a volatile anesthetic.

Other targets in addition to Na_v_ likely contribute to the neuronal anesthetic actions, and provide additional possibilities for synaptic diversity and thus pharmacological selectivity. Interneuron diversity derives in part from differences in synaptic signaling protein expression (40). Several types of presynaptic proteins are affected by clinical concentrations of isoflurane, including ion channels, Ca^2+^-binding proteins, and intracellular signaling proteins (41). Other potential targets for transmitter-selective or interneuron-selective actions include K^+^ channels, which determine neuronal excitability, AP morphology, and ultimately neurotransmitter release via membrane hyperpolarization and repolarization (42), mitochondrial electron transport complex proteins involved in presynaptic energetics (43), and presynaptic SV exocytotic proteins such as syntaxin (44).

### Differential Na_v_ subtype expression and pharmacological consequences

Hippocampal and cortical pyramidal glutamatergic neurons express Na_v_1.1 mostly in their somata and dendrites, while Na_v_1.6 is expressed primarily in axons and the axonal initial segment where it controls axonal excitability. Interneurons express more Na_v_1.1, with PV^+^ interneurons dependent on axonal Na_v_1.1 for their unique firing properties (21). The importance of Na_v_1.1 to interneuron physiology is exemplified in Dravet’s syndrome, an Na_v_1.1 channelopathy that causes PV^+^ neuron dysfunction while sparing pyramidal neurons (45). Although glutamatergic neurons also express Na_v_1.1, lower levels of expression and different subcellular localization appear to limit the consequences of Na_v_1.1 loss-of-function on signaling and hyperexcitability. Knockdown of Na_v_1.1 in glutamatergic neurons did not affect the sensitivity of SV exocytosis to isoflurane, suggesting that Na_v_1.1 contributes less to SV exocytosis in glutamatergic than in GABAergic neurons due to its reduced expression or axonal enrichment.

Expression of Na_v_ differs between cortical and hippocampal cell types, with PV^+^ and SST^+^ interneurons differing in both Na_v_ amount and subcellular expression. In PV^+^ basket cells, Na_v_ is enriched in axons and nearly absent in dendrites (6), whereas SST^+^ interneurons have enriched somatodendritic Na_v_ (46, 47). A recent genome-wide association study of cortical interneuron subtype expression patterns based on RNAseq showed that PV^+^ neurons express more Na_v_1.1 mRNA than SST^+^ neurons (22). While SST^+^ interneuron axons express more Na_v_1.2 and Na_v_1.2-mediated axonal currents, neither are observed in PV^+^ interneurons (25), suggesting that SST^+^ neurons rely more on Na_v_1.2 than Na_v_1.1 to mediate axonal excitability. Hippocampal RNAseq also confirms that PV^+^ interneurons contain more Na_v_1.1 transcript than SST^+^ interneurons, which have more Na_v_1.2 (48). Sensitivity of Na_v_1.2 to isoflurane is similar to that of Na_v_1.6. Therefore, differential Na_v_ expression underlies synapse diversity and complexity that has pharmacological implications for anesthetic actions on SV exocytosis from PV^+^ *versus* SST^+^ interneurons.

The role of Na_v_1.1 in determining the neurophysiological and pharmacological properties of interneuron subtypes is shown by our observation that Na_v_1.1 overexpression in SST^+^ neurons reduced the potency of isoflurane for inhibition of SV exocytosis. Although SST^+^ interneurons express more Na_v_1.1 than glutamatergic neurons, we found similar sensitivity to isoflurane inhibition of SV exocytosis. Thus SST^+^ Na_v_1.1 may have a minimal role in anesthetic actions of SV exocytosis due to greater dendritic than axonal trafficking. It is also possible that SST^+^ neurons express another important anesthetic-sensitive target (*vida infra*). Because our experiments were performed using TTX to block endogenous Na_v_-mediated exocytosis, we propose that heterologous expression of Na_v_1.1 in axons supersedes the native dendritic expression. We propose that SST^+^ neurons are more sensitive than PV^+^ neurons to anesthetic effects on SV exocytosis due to differences in the relative expression of Na_v_1.1 *versus* Na_v_1.2 and Na_v_1.6, particularly in presynaptic sites that control nerve terminal excitability.

Anesthetic inhibition of Na_v_ could disrupt AP generation at the axon initial segment, conduction at the nodes of Ranvier, and/or entry into and depolarization of the presynaptic terminal. Recent advances in high-sensitivity optical voltage indicators could allow direct compartment-specific quantification of measures of anesthetic actions on presynaptic APs from neurons with differences in expression of specifc Na_v_ subtypes(49).

### Implications for neuronal network function

Interneurons are central modulators of neuronal excitability and network function, with cell-specific roles in organizing network activity and driving oscillatory rhythms (50). Anesthesia is associated with major alterations in macroscopic network oscillations that are thought to be important to loss of consciousness (33). Thus, understanding cell-specific anesthetic actions on interneuron output is necessary to understand the neurophysiology of the anesthetized state, including the population electrical patterns and oscillations that characterize distinct stages of sedation and hypnosis. Cortical EEG patterns are routinely used intra-operatively to monitor anesthetic depth, and hippocampal interneuron cell-types have cortical homologues with comparable output properties (5). PV^+^ interneurons are instrumental in generating gamma oscillations, so their behavior under anesthesia is especially important to understand(31). Since interneurons control neuronal excitation, these effects are also broadly relevant to experiments performed under isoflurane anesthesia, such as *in vivo* electrophysiological or optical recordings.

How behavioral and macroscopic features such as disruption of normal oscillatory rhythms and unconsciousness emerge from anesthetic actions on synaptic function remains unclear, in part because synaptic actions are more complex than a global imbalance of excitation/inhibition, and cortical and hippocampal interneurons play distinct roles in neural circuits. Thus, determining interneuron subtype-specific anesthetic effects is critical to understanding their network level actions. Single cell transcriptome analysis will likely prove invaluable in investigating cell and circuit specific pharmacology, and in identifying novel drug targets that selectively impair the critical CNS circuits for consciousness and pain.

### Limitations

Neuron-specific effects on the action potential or presynaptic excitability due to Na_v_ subtype-specific pharmacology can explain the observed effects of Na_v_ manipulation on anesthetic sensitivity. However direct measurement of presynaptic voltage and/or Na^+^ currents, which has hitherto been experimentally intractable, is necessary to confirm this mechanism.

Interneuron subtype diversity has been studied intensely and catalogued in intact mouse brain, however it is unclear if normal developmental gene expression patterns and connectivity profiles are recapitulated in primary dissociated neuron cultures. Although *in vitro* neuronal cultures provide a good model for investigating the molecular and cellular mechanisms of synaptic transmission, they are limited as a proxy for intact neuronal networks. Expression of PV is developmentally delayed, beginning at DIV8 and plateauing at DIV28. Since the optimal age for syn-pH based SV exocytosis recording is <DIV21, developing PV interneurons weakly express PV at the time of recording and may be misidentified as PV^-^. However, our use of retrospective immunostaining for PV protein biases our recordings for PV^+^ neurons advanced toward the mature phenotype.

Given the lack of Na_v_ subtype-selective antagonists, we employed TTX-resistant Na_v_ mutants to probe the function of specific subtypes in intact neurons. Overexpression of transfected Na_v_ could lead to ectopic expression in cellular compartments that are not normally occupied by native Na_v_. There is no evidence that Na_v_1.1_R_ and Na_v_1.6 _R_ have different pharmacological properties than wild-type channels, as both are rendered TTX-resistant by single point mutations in the extracellular toxin binding domain, and anesthetic sensitivities and gating properties are unchanged (23, 24).

Na_v_1.1 loss of function is associated with aberrant interneuron function and an epileptic phenotype, so compromised cell health due to reduced Na_v_1.1 is possible. We interpreted the ineffectiveness of Na_v_1.1 knockdown on anesthetic inhibition of SV exocytosis in glutamatergic neurons as indicating a lesser role of Na_v_1.1 in terminal excitability. However it is possible that knockdown was incomplete and insufficient to affect exocytosis.

## Conclusions

We identified a role for differential Na_v_ subtype expression in synaptic diversity resulting in pharmacological selectivity in the effects of the volatile anesthetic isoflurane on SV exocytosis in glutamatergic and GABAergic hippocampal neurons This provides a potential mechanistic link between anesthetic effects on channel function and subtype-specific effects on presynaptic terminal depolarization, Ca^2+^ entry, and neurotransmitter release. The anesthetic-resistant Na_v_1.1 subtype plays a prominent functional role in PV^+^ interneurons, and we hypothesize that the PV^+^ action potential is relatively resistant to anesthetic depression. In contrast, greater expression of the anesthetic-sensitive Na_v_1.2 and Na_v_1.6 subtypes in glutamatergic and SST^+^ interneurons leads to reduced activity-dependent SV exocytosis, potentially due to greater action potential depression. Further studies are required to determine changes in presynaptic Ca^2+^ and membrane potential in identified interneuron subtypes. Neuronal subtype-selective anesthetic effects may link the molecular mechanisms of anesthetic action to higher-order systems-level and behavioral endpoints of anesthesia, which remains one of the most important gaps in our conceptual understanding of general anesthesia.

## Methods

### Mice

Postnatal GAD2-Cre::Ai14-tdTomato, SST-Cre::Ai14-tdTomato, and Nkx2.1-Cre::Ai14 mice were generated by breeding GAD2-Cre, SST-Cre, and Nkx2.1-Cre mice singly to Ai14 mice. All mice were commercially obtained from Jackson Laboratories (Bar Harbor, Maine; stock #010802, #013044, #008661, and #007914). All experimental procedures were approved by the Weill Cornell Medical College Institutional Animal Care and Use Committee and conformed to NIH Guidelines for the Care and Use of Animals.

### Neuron culture and transfection

Hippocampi were dissected from postnatal mice (0-2 days old, both sexes). Cells were dissociated and plated onto polyornithine-coated coverslips as described (39). Transfection was performed on day 6 or 7 *in vitro* (DIV6-7) using a calcium phosphate-mediated gene transfer protocol for sparse transfection (39) experimentally adjusted to yield ∼0.75 transfected GAD2+ neurons per coverslip. For each experiment, neurons were derived from at least three separate primary hippocampal neuron preparations.

### Plasmids

Synaptophysin-pHluorin2x (syn-pH), a chimera of synaptophysin with two copies of pHluorin (2x) inserted into the second intravesicular loop, was used as an optical biosensor for SV exocytosis (51). For Na_v_ overexpression studies, syn-pH was co-transfected with plasmids encoding TTX-resistant mutants for human Na_v_1.1_R_ [hNa_v_1.1F383S] or mouse Na_v_1.6_R_ [mNa_v_1.6Y371S] (37-39). For knockdown of endogenous Na_v_1.1, syn-pH was co-transfected with a validated shRNA plasmid against the mRNA target sequence TGCCAATGCATGTCGATATTA in the mouse Nav1.1 5’ UTR.

### Live-cell Imaging of SV Exocytosis

Experiments were performed on a ZEISS Axio Observer Z1 widefield fluorescence microscope (Oberkochen, Germany). Fluorescence was imaged with an Andor iXon1 EMCCD (Belfast, Ireland), filter cubes for eGFP and RFP and LED illumination (Colibri 7 System, Zeiss). Coverslips were mounted in a closed-bath field stimulation perfusion chamber with a total volume of 263 μl; live-cell imaging was performed at 37.0 ± 0.2°C, with temperature maintained by an objective heater, in-line solution heater, and imaging chamber heater (Warner Instruments, Hamden, CT). Solutions were continuously perfused at 1ml/min using a custom system. The standard buffer was Tyrode’s solution (119 mM NaCl, 2.5 mM KCl, 2 mM CaCl_2_, 2mM MgCl_2_, 25 mM HEPES buffered to pH 7.4, 30 mM glucose) containing 10 μM 6-cyano-7-nitroquinoxaline-2,3-dione (CNQX) and 50 μM D,L-2-amino-5-phosphonovaleric acid (AP5) to block glutamatergic signaling (both from Tocris, Bristol, UK). Experiments involving tetrodotoxin (TTX) resistant Na_v_ mutants were performed in 4 mM Ca^2+^ Tyrode’s solution, modified to 4 mM CaCl_2_ and 0 mM MgCl_2_, plus 300 nM TTX diluted from 1 mM stock aliquots (Alomone, Jersalem, Israel). Imaging solutions were perfused using a multi-barrel manifold (Automate Scientific, Berkeley, CA).

Transfected neurons were identified by resting syn-pH fluorescence in boutons and axons, and cell-type was identified based on the presence or absence of overlapping tdTomato/synpH observed in a fluorescence overlay. To drive exocytosis, action potential (AP) trains were generated by field stimulation with a pulse generator (Master-9, A.M.P.I., Jersalem, Israel) and platinum/iridium bath electrodes built into the imaging chamber (10 V/cm^2^).

An experimental solution of 0.5 mM isoflurane dissolved in Tyrode’s solution was prepared daily from 12 mM saturated stock solutions of isoflurane, and focally perfused from gas-tight glass syringes onto neurons in the closed-bath imaging chamber through the injection port. This clinically relevant concentration corresponds to ∼1.4 MAC (minimum alveolar concentration) in mice (40). Isoflurane was applied for 5 min before imaging to allow equilibration. At the conclusion of each experiment, a perfusate sample was taken from the chamber for analysis of delivered isoflurane concentration using a Shimadzu GC-2010 Plus gas chromatograph (Kyoto, Japan) with external standard calibration (52). The reported value of 0.50±0.039 mM reflects the averaged measured from all bath samples collected. Total pool of SVs was determined after the washout recording by perfusing 50 mM NH_4_Cl (substituted for 50 mM NaCl in Tyrode’s solution, buffered to pH 7.4) to alkalize vesicles and reveal total vesicular syn-pH.

### shRNA Validation

Pre-validated shRNA plasmids were identified on the Broad Institute RNAi Consortium shRNA Library Database and purchased from Sigma-Aldrich (St. Louis, MO). Multiple shRNA plasmids were tested by immunostain analysis of Na_v_1.1 density (see **Supplemental Figure 5** for validation data). Quantitative Na_v_1.1 immunostaining analysis was performed using published protocols (53). Based on these preliminary experiments, clone TRCN0000225834 was further tested by rescue with Na_v_1.1_R_.

### Immunocytochemistry

Cultured neurons were fixed with 3.7% formaldehyde and 4% sucrose in PBS for 15 min (150 mM NaCl, 2.7 mM KCl, 10 mM Na_2_HPO_4_, 2 mM KH_2_PO_4_, pH 7.4), permeabilized with 0.2% Triton X-100 for 30 min, blocked in 5% bovine serum albumin in PBS for 30 min, and incubated with appropriate primary antibody diluted in 1% BSA for 4°C overnight. Primary antibodies were against parvalbumin (rabbit, Abcam 11427, Cambridge, UK), Na_v_1.1, and Na_v_1.6 (rabbit, Alomone ASC-001 and ASC-008, Jersalem, Israel). Neurons immunolabelled with an anti-Na_v_1.1 primary antibody were detected with anti-rabbit Alexa-488-conjugated goat secondary antibody, and neurons immunolabeled with an anti-parvalbumin primary antibody were detected with anti-rabbit Alexa-633-conjugated secondary antibody (1:1000, in 1% BSA/PBS). After a four-hour incubation with secondary antibodies, coverslips were mounted and imaged with a confocal microscope. Alexa Fluor 488- and 633-conjugated antibodies were obtained from Life Technologies (Carlsbad, CA).

For optical exocytosis measurements in Nkx2.1-Cre::Ai14 cultures, retrospective immunolabelling was used to identify PV^+^ interneurons following live-cell imaging (26). After recording SV exocytosis, coverslips were removed from the imaging chamber, and neurons were fixed as above and stored at 4°C for up to two weeks. Immunostaining was performed as above using rabbit anti-parvalbumin monoclonal antibody (ab11427, Abcam, Cambridge, UK) and goat anti-rabbit Alexa Fluor 633-conjugated secondary antibody. Recorded syn-pH-expressing boutons were located on the coverslip by stage coordinates, and analyzed for PV immunoreactivity.

### Image and statistical analysis

Live-cell imaging recordings were analyzed in ImageJ (rsb.info.nih.gov/ij) using a custom plugin (rsb.info.nih.gov/ij/plugins/time-series.html) to measure fluorescence over time within 2 μm diameter regions of interest (ROIs), with each ROI containing a synaptic bouton. Transfected boutons were selected as ROIs based on their response to control stimulation of 100 APs in Tyrode’s buffer. ROI cell-type was identified by colocalization of RFP with GFP (resting syn-pH fluorescence). A washout measure significantly smaller than control indicates deteriorating cell condition, and cells with washout response < 60% of control were excluded from analysis

Candidate ROIs were identified using the maximal exocytosis fluorescence signal (FMax) and verified by calculating evoked changes in fluorescence. Isoflurane effects were calculated for each ROI. First, fluorescence was normalized to the syn-pH total vesicular pool, or the maximum intensity with NH_4_Cl alkalization. Normalized values were used to calculate evoked SV exocytosis signal (ΔF = [FMax-F0]/F0) for each treatment, and the isoflurane effect (ΔI= [IsofluraneΔF - ControlΔF]/ControlΔF). For each bouton, signal:noise ratio (SNR) was defined as SNR=ΔF/σ, where ΔF is [FMax-F0]/F0 and s is the standard deviation (SD) of the fluorescence measured for 10 frames before stimulation. Boutons with SNR <4 were excluded from analysis. Remaining ROIs were pooled to create an ensemble single-cell average. Values are shown as mean (SD). ANOVA with Tukey’s *post hoc* test, Student’s or Welch’s unpaired t test (P < 0.05), and 95% confidence intervals on the difference of the means were used to determine statistical significance, defined as P>0.05.

## ACKNOWLEDGMENTS

We thank members of the Hemmings laboratory for constructive comments. We also thank Jaime de Juan Sanz for guidance in shRNA experiments. This work was supported by National Institutes of Health Grant R01 GM58055 (to H.C.H.).

## Supplemental Figures

**Figure S1: Repeated SV exocytosis measured from putative glutamatergic and GABAergic neurons in GAD2-Cre::Ai14 hippocampal neurons**. Connected dot plots show individual cell peak FΔ/F_0_ values from glutamatergic (**A**, GAD2^−^) and GABAergic (**B**, GAD2^+^) neurons following control, isoflurane, and washout treatments. Isoflurane was omitted for a subset of recordings (- isoflurane) to observe the effect of repeated electrical stimulation on SV exocytosis.

**Figure S2: Distribution of GAD2**^**+**^ **and GAD2**^−^ **responses to isoflurane inhibition of SV exocytosis**. Responses were measured following transfection of GAD2-Cre::Ai14 transgenic mouse hippocampal neurons with syn-pH, and binned by per cent inhibition relative to control (no isoflurane) as per cent of total boutons with that response for each neuron type. n=24 GAD2^+^ neurons and n=13 GAD2^-^ neurons.

**Figure S3: Synaptic expression of parvalbumin and syn-pH in Nkx2**.**1-Cre::Ai14 cultures**. Parvalbumin, a cytosolic protein present in both the soma and presynaptic terminal, is a marker for a subclass of GABAergic interneurons that can be identified by retrospective immunocytochemical analysis following syn-pH SV exocytosis measurements. Hippocampal neurons from Nkx2.1-Cre::Ai14 mice expressing tdTomato and transfected with syn-pH. **A**. String of syn-pH expressing boutons that are not immunoreactive for PV (arrowheads). **B**. String of syn-pH expressing boutons immunoreactive for PV (arrowheads). **C**. Higher magnification images showing syn-pH^+^ boutons that are PV^+^ and those that are PV^−^ within the same field of view (+ and – arrowheads, respectively).

**Figure S4: Validation of Na**_**v**_**1**.**1**_**R**_ **and Na**_**v**_**1**.**6**_**R**_ **overexpression. A**. Representative images of native Na_**v**_1.1 and Na_**v**_1.6 immunoreactivity in cultured wild-type mouse hippocampal neurons. Scale bars represent 10 µm. **B**. Na_**v**_1.1_R_ and **C**. Na_**v**_1.6_R_ overexpression was quantified by immunocytochemistry comparing wild-type control neurons transfected with cytosolic mCherry alone with neurons co-transfected with mCherry and hNa_v_1.1_R_ or mNa_v_1.6_R_. Total Na_v_1.1 or 1.6 immunoreactivity within transfected processes, representing both native and transfected Na_v_, was quantified using threshold-based masks drawn around proximal axons using the mCherry signal. Na_v_ immunofluorescence (green) within the mask was normalized to mCherry fluorescence (red), both quantified as raw integrated density (the sum of all the pixel intensities in the ROI) using ImageJ. Na_v_1.1_R_ or Na_v_1.6_R_ Int. Density is presented after normalization to mCherry. Na_v_1.1_R_ *vs*. control, p=0.0046; Na_v_1.6_R_ *vs*. control, p=0.003 (t-tests). Scale bars represent 10 µm.

**Figure S5. Validation of Na**_**v**_**1**.**1 knockdown and overexpression rescue by immunocytochemistry. A**. To quantify shRNA knockdown efficiency, wild-type mouse hippocampal neurons were transfected with mCherry (control), or mCherry plus an shRNA construct targeted to the Na_v_1.1 3’UTR (ID: TRCN0000225834, Sigma-Aldrich). In control neurons strong Na_v_1.1 immunoreactivity (green) was observed in transfected and in adjacent untransfected cells. To quantify Na_v_1.1 density within transfected processes, threshold-based masks were drawn around a proximal axon using the mCherry signal (white outline). The Na_v_1.1 signal within the mask was quantified as an integrated density normalized to mCherry integrated density. **B**. Comparison of control, shRNA, and shRNA rescued with hNa_v_1.1_R_. Mean shRNA knockdown efficiency was 57% of control. Because the native mouse Na_v_1.1 3’UTR is not present in the hNa_v_1.1_R_ expression construct, the mutant protein resists knockdown. Control *vs* shRNA, p=0.0066; shRNA *vs*. shRNA + Nav1.1_R_, p=0.052; control *vs*. shRNA + Nav1.1_R_ p=0.0836; ANOVA with Tukey’s *post hoc* test.

