## Supplementary figures and images for "Differential inhibition of GABA release from mouse hippocampal interneuron subtypes by the volatile anesthetic isoflurane"

### Supplemental Figure 1

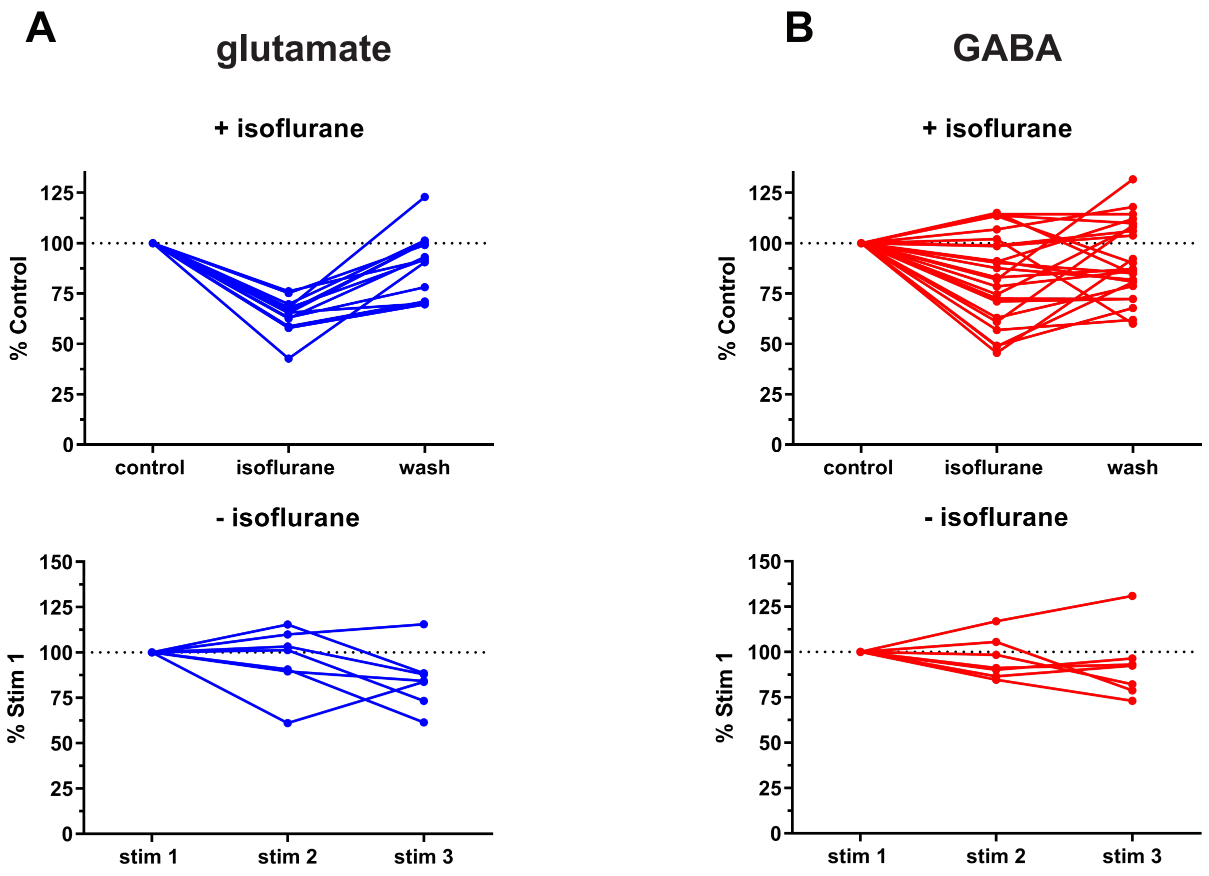

### Supplemental Figure 2

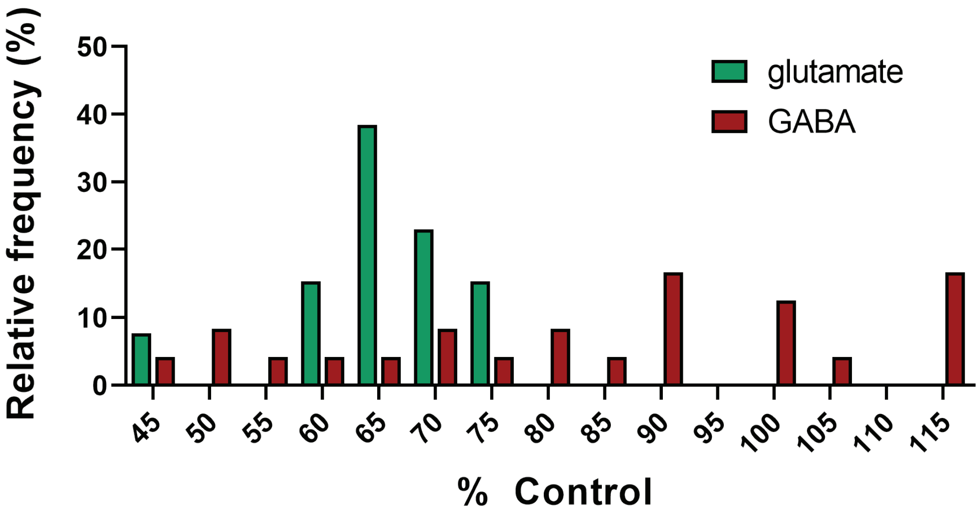

### Supplemental Figure 3

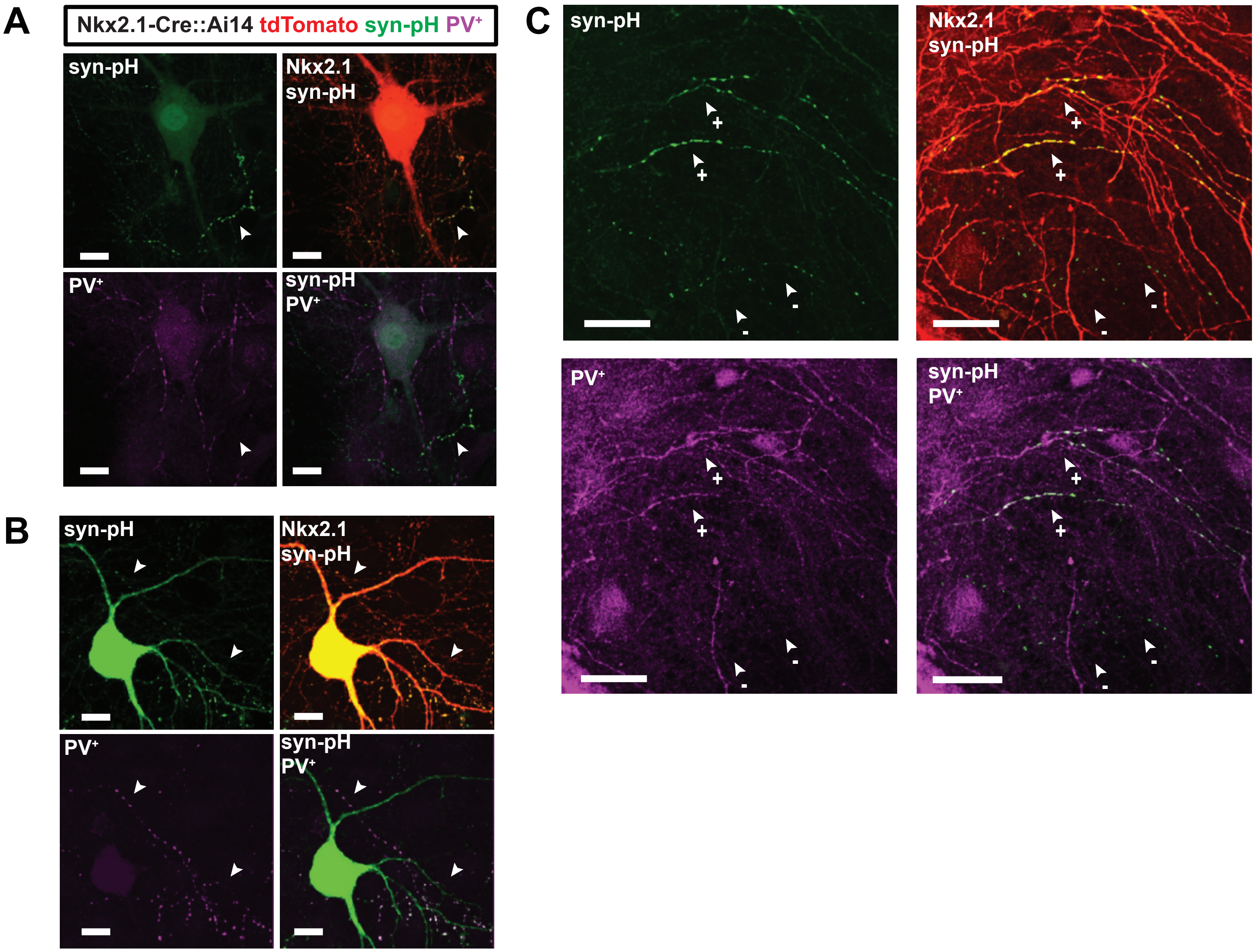

### Supplemental Figure 4

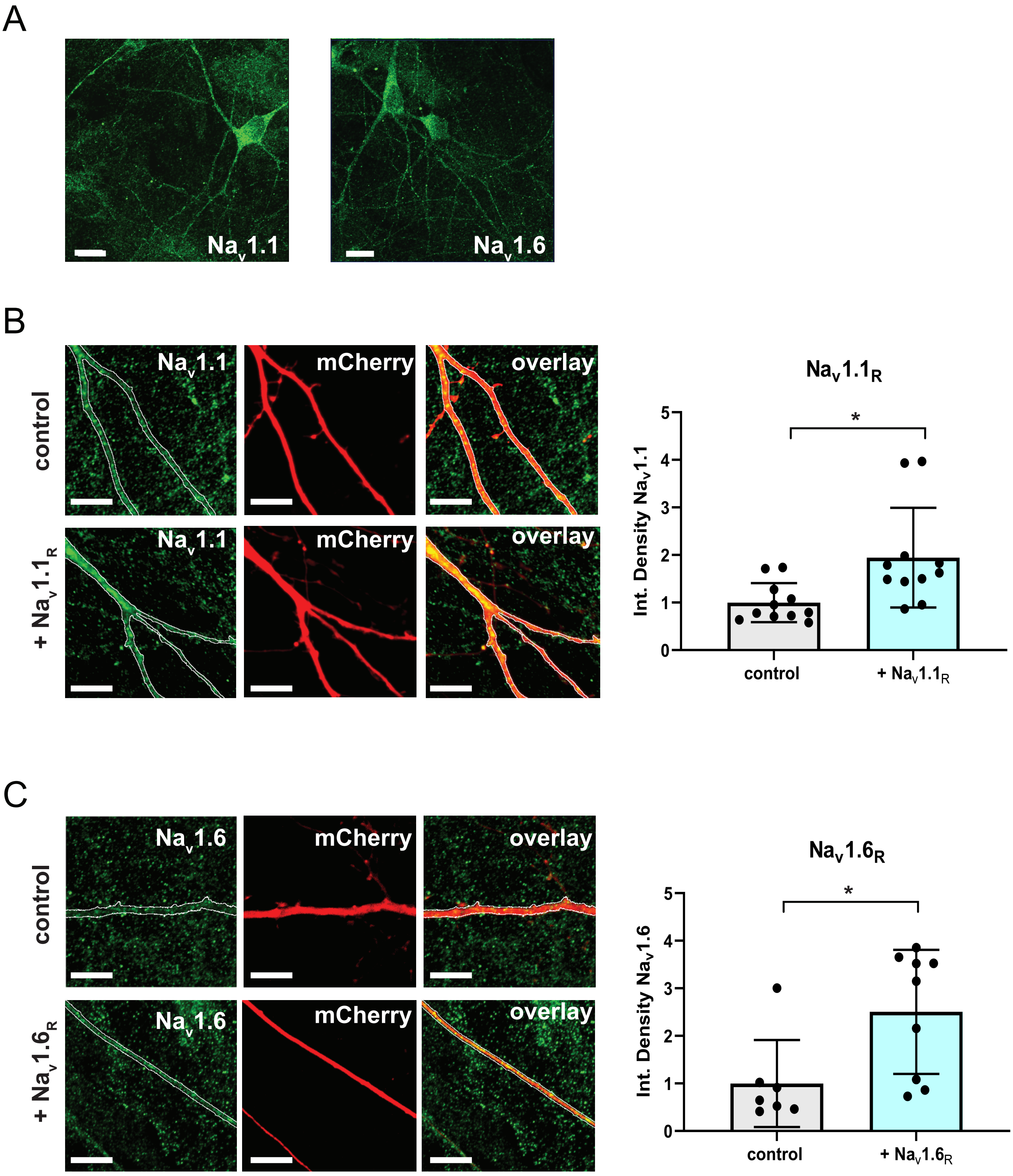

### Supplemental Figure 5

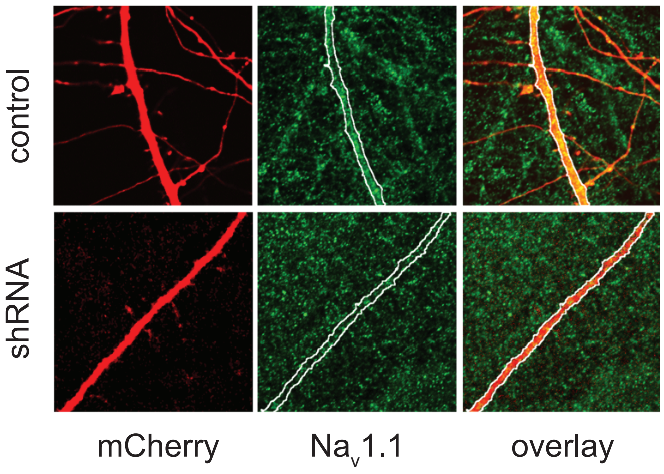

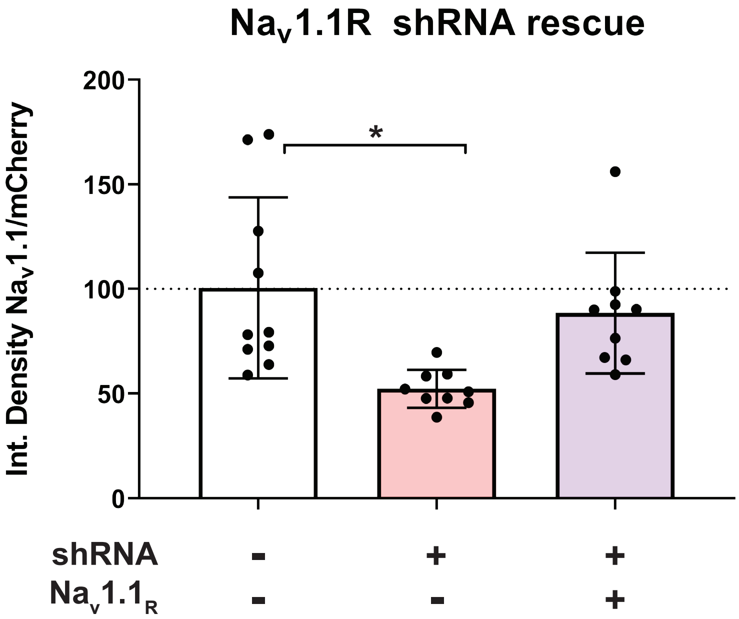
